# Population coding and oscillatory subspace synchronization integrate context into actions

**DOI:** 10.1101/2021.12.17.473118

**Authors:** Jan Weber, Anne-Kristin Solbakk, Alejandro O. Blenkmann, Anais Llorens, Ingrid Funderud, Sabine Leske, Pål Gunnar Larsson, Jugoslav Ivanovic, Robert T. Knight, Tor Endestad, Randolph F. Helfrich

**Affiliations:** Hertie Institute for Clinical Brain Research, Center for Neurology, University Medical Center Tübingen, Tübingen, Germany; International Max Planck Research School for the Mechanisms of Mental Function and Dysfunction, University of Tübingen, Tübingen, Germany; Department of Psychology, University of Oslo, Oslo, Norway; RITMO Centre for Interdisciplinary Studies in Rhythm, Time and Motion, University of Oslo, Oslo, Norway; Department of Neurosurgery, Oslo University Hospital, Oslo, Norway; Department of Neuropsychology, Helgeland Hospital, Mosjøen, Norway; Helen Wills Neuroscience Institute, UC Berkeley, Berkeley, CA, USA; Department of Musicology, University of Oslo, Oslo, Norway; Department of Psychology, UC Berkeley, Berkeley, CA, USA

## Abstract

Contextual cues and prior evidence guide human goal-directed behavior. To date, the neurophysiological mechanisms that implement contextual priors to guide subsequent actions remain unclear. Here we demonstrate that increasing behavioral uncertainty introduces a shift from an oscillatory to a continuous processing mode in human prefrontal cortex. At the population level, we found that oscillatory and continuous dynamics reflect dissociable signatures that support distinct aspects of encoding, transmission and execution of context-dependent action plans. We show that prefrontal population activity encodes predictive context and action plans in serially unfolding orthogonal subspaces, while prefrontal-motor theta oscillations synchronize action-encoding population subspaces to mediate the hand-off of action plans. Collectively, our results reveal how two key features of large-scale population activity, namely continuous population trajectories and oscillatory synchrony, operate in concert to guide context-dependent human behavior.

## Introduction

Human decisions strongly depend on available prior evidence and contextual cues. A long-standing question in models of top-down guided behavior is how prior evidence is incorporated into momentary action plans^1-3^. The active sensing framework postulates that the brain utilizes its inherent rhythmic structure as an energy-efficient mechanism to implement temporal predictions^4,5^. This framework further predicts that the brain switches from a rhythmic to a continuous, and therefore energy-costly, processing mode when less prior evidence is available. Furthermore, active sensing implies that synchronization of endogenous oscillations is instrumental for inter-areal information transfer, as suggested by the communication-through-coherence hypothesis^6^. Active sensing has mainly been studied in the context of sensory selection^7-9^ and to date it remains unknown whether similar principles apply when context is signaled by abstract cues. In a different line of research, recent work in non-human primates (NHP) has demonstrated that sensorimotor cortex as well as adjacent premotor areas, such as the frontal eye fields, encode high-level contextual information in neural population codes^10-15^. Whereas the active sensing framework relies on univariate features (i.e., oscillatory power, phase, and neural firing), the population doctrine emphasizes that information is encoded in the entire population response that can be conceptualized as a trajectory passing through a high-dimensional neural state space^16^. To date, population coding and neural oscillations, two key signatures of coordinated population activity, have mainly been studied in isolation. Consequently, it remains elusive how both features interact to guide goal-directed behavior.

In this study, we addressed how high-level contextual information is flexibly integrated into current action plans in humans. We specifically tested if principles of the active sensing framework also apply to prefrontal-motor interactions when contextual information is rule-based and not sensory-driven^9^. Furthermore, we aimed to determine the population correlates of presumed rhythmic and continuous processing modes. So far, population activity has mainly been studied using single- and multi-unit recordings in NHP. Here we recorded intracranial electroencephalography (iEEG) from prefrontal and motor cortex in patents with epilepsy who underwent invasive monitoring for localization of the seizure onset zone. We specifically studied high-frequency band activity (HFA; 70-150 Hz) as a proxy of population firing to address if coding principles that have previously been identified in NHP also apply in the human brain. All participants engaged in a predictive motor task, where they were instructed to continuously track a moving target and release a button once it reached a pre-defined spatial location. A contextual cue determined the probability of a premature and abrupt stop when participants had to withhold their ongoing response.

We describe a striking functional dissociation between population activity and network oscillations where human PFC encodes predictive context and the current action plan in orthogonal subspaces using a continuous processing regime, while theta oscillations mediate the hand-off of the current action plan from prefrontal to motor regions. Collectively, we identified computationally distinct roles of continuous and rhythmic brain activity at the population level that jointly guide context-dependent, goal-directed human behavior.

## Results

We recorded intracranial EEG (iEEG) from 19 pharmaco-resistant patients with epilepsy (33.73 years ± 12.52, mean ± SD; 7 females) who performed a predictive motor task (**Fig. 1a**). Participants had to closely track a moving target and respond (go trial) as soon as the target reached a predefined spatial location (hit lower limit; HLL). They were instructed to withhold their response if the target stopped prematurely (stop trial). A predictive cue indicated the likelihood of a stop trial (green circle = 0%, orange circle = 25%, red circle = 75%). We refer to the stop likelihood as behavioral uncertainty or predictive context and use these terms interchangeably. We simultaneously recorded from prefrontal cortex (PFC) and motor cortex to study how the human prefrontal-motor network converts predictive context into concrete actions (**Fig. 1b**).

**Fig. 1.**
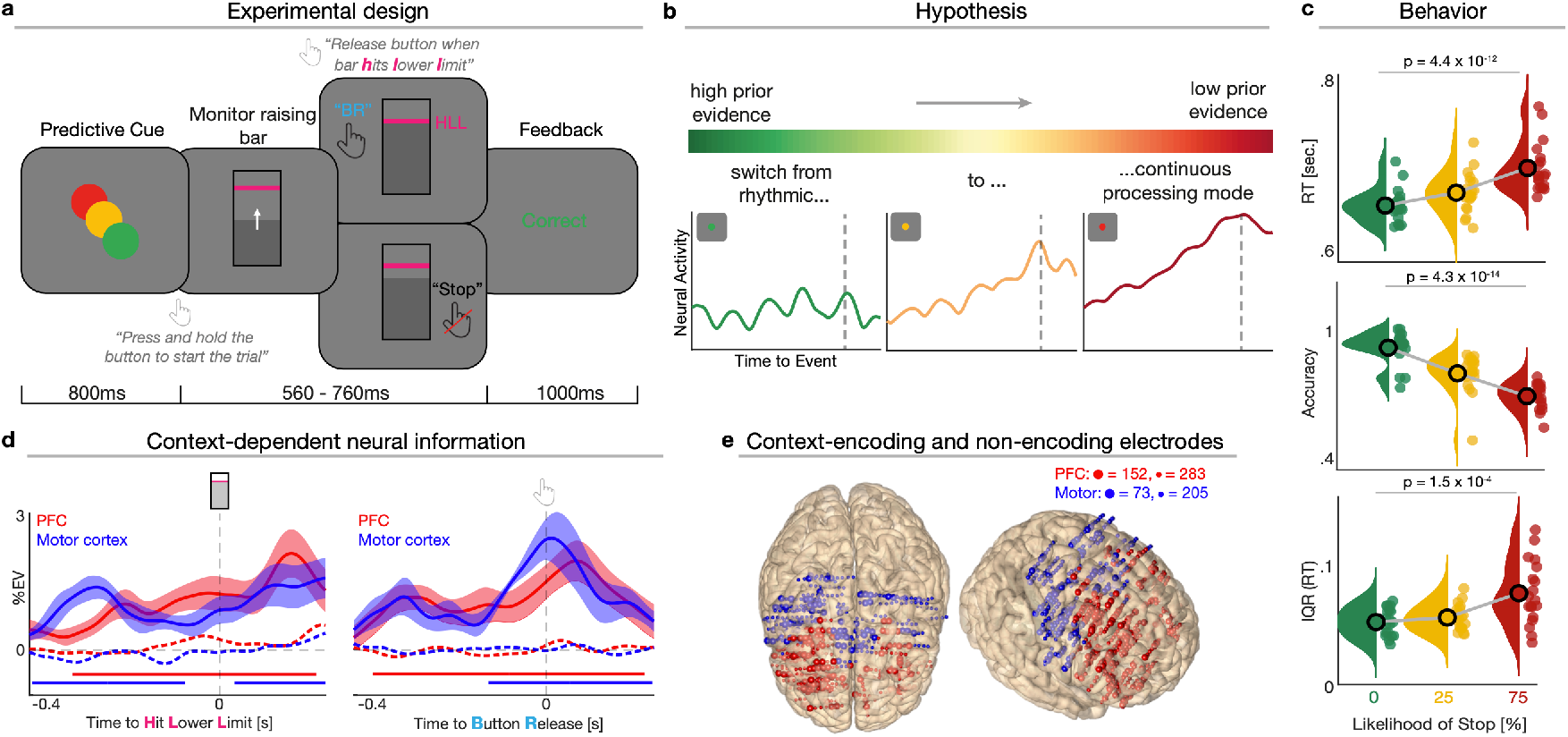
Task design, hypothesis, behavioral results and electrophysiological signatures of context-dependent neural information. **a**, Participants were presented with a predictive cue indicating the likelihood that a moving target (self-initiated via space bar press) would stop prior to a predefined lower limit (HLL; pink horizontal line). Participants were asked to release the space bar as soon as the target hit the lower limit (go trial) or withhold the response if the target stopped before reaching the lower limit (stop trial). Afterwards, participants received feedback. **b**, Schematic illustration of our key hypothesis. States of high behavioral uncertainty should introduce a switch towards stronger ramping dynamics. **c**, Behavioral results. Scattered dots represent single grand averages, black outlined dots depict the group level average and histograms illustrate the probability distribution. Upper: RTs gradually scales with behavioral uncertainty. Middle: Accuracy gradually decreases as a function of behavioral uncertainty. Lower: Interquartile range also increases from trials with no to high behavioral uncertainty. **d**, ROI-specific time course of context-dependent neural information (percent explained variance, %EV) for context-encoding (solid lines) and non-encoding electrodes (dashed lines). The lower horizontal lines show the temporal extent of significant cluster differences between context-encoding and non-encoding electrodes for the respective ROI. Shading represents the standard error of the mean (SEM) across participants. **e**, Context-encoding (large spheres) and non-encoding (small spheres) electrodes overlaid on a standardized brain in MNI space for our two regions of interest.

### Neural and behavioral signatures of context-dependent evidence accumulation

We confirmed that participants used the predictive cue to guide behavior. We found that reaction times (RT) gradually increased as a function of uncertainty (**Fig. 1c**; +43.9ms ± 19.8ms, mean ± SD; *F*_2,36_ = 58.99, *P* = 4.4 × 10^−12^, 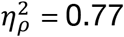; one-way RM-ANOVA). Participants were also significantly less accurate in trials with high uncertainty (**Fig. 1c**; -20.7% ± 8%; mean ± SD; *F*_2,36_ = 81.53, *P* = 4.3 × 10^−14^, 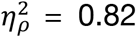). Furthermore, we quantified how predictive context modulated participants’ sensitivity *d*’ (d-prime) and decision criterion *c* (Methods). We found that *d*’ decreased (**Supplementary Fig. 1**; *P* = 0.002, Cohen’s *d* = 1.03; Wilcoxon rank sum test) and *c* increased (**Supplementary Fig. 1**; *P* = 0.008, Cohen’s *d* = -0.7) with uncertainty. To quantify trial-by-trial variability, we assessed the interquartile range (IQR) as a measure of dispersion (**Fig. 1c**). We found that RTs were more consistent for predictive trials (IQR 0.05s ± 0.01s; mean ± SD) and more variable under high uncertainty (IQR 0.08s ± 0.03s; mean ± SD; *F*_2,36_ = 11.36, *P* = 1.5 × 10^−4^, 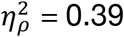; one-way RM-ANOVA). In sum, these results demonstrate that states of high behavioral uncertainty are detrimental for the speed, accuracy, and sensitivity of action-linked decisions.

We subsequently determined the neural dynamics that underlie behavioral uncertainty by assessing HFA as a proxy for local population activity^17-19^. The initial quantification of percent variance^20-24^ explained by context revealed significant context-dependent neural information in both PFC and motor cortex when time-locked to the HLL (**Fig. 1d**; *PFC*: t_15_ = 985.91, *P* < 0.001, Cohen’s *d* = 0.83; *motor cortex*: first cluster, t_10_ = 761.78, *P* < 0.001, Cohen’s *d* = 1.24; second cluster, t_10_ = 351.6, *P* < 0.001, Cohen’s *d* = 0.96; cluster test). A comparable pattern was observed when time-locked to action execution (button release, BR; *PFC*: t_16_ = 1144.2, *P* < 0.001, Cohen’s *d* = 0.88; *motor cortex*: t_10_ = 941.25, *P* = 0.0016, Cohen’s *d* = 1.51). Neural information evolved similarly in both regions over time (all *P* > 0.09; cluster test).

Overall, we found that 35% (N = 152) of all electrodes in PFC and 27% (N = 73) of all electrodes in motor cortex significantly encoded context (**Fig. 1e**; Methods). We used context-encoding electrodes for subsequent univariate analyses unless stated otherwise. We found a context-dependent HFA modulation in both *PFC* (**Fig. 2a**; first cluster: *F*_2,30_ = 699.15, *P* = 0.009; second cluster: *F*_2,30_ = 496.57, *P* = 0.018) and *motor cortex* (**Fig. 2b**; *F*_2,20_ = 326.6, *P* = 0.036). The strongest context-dependent modulation was observed for the PFC ~300ms prior to the HLL (**Fig. 2a)**.

**Fig. 2.**
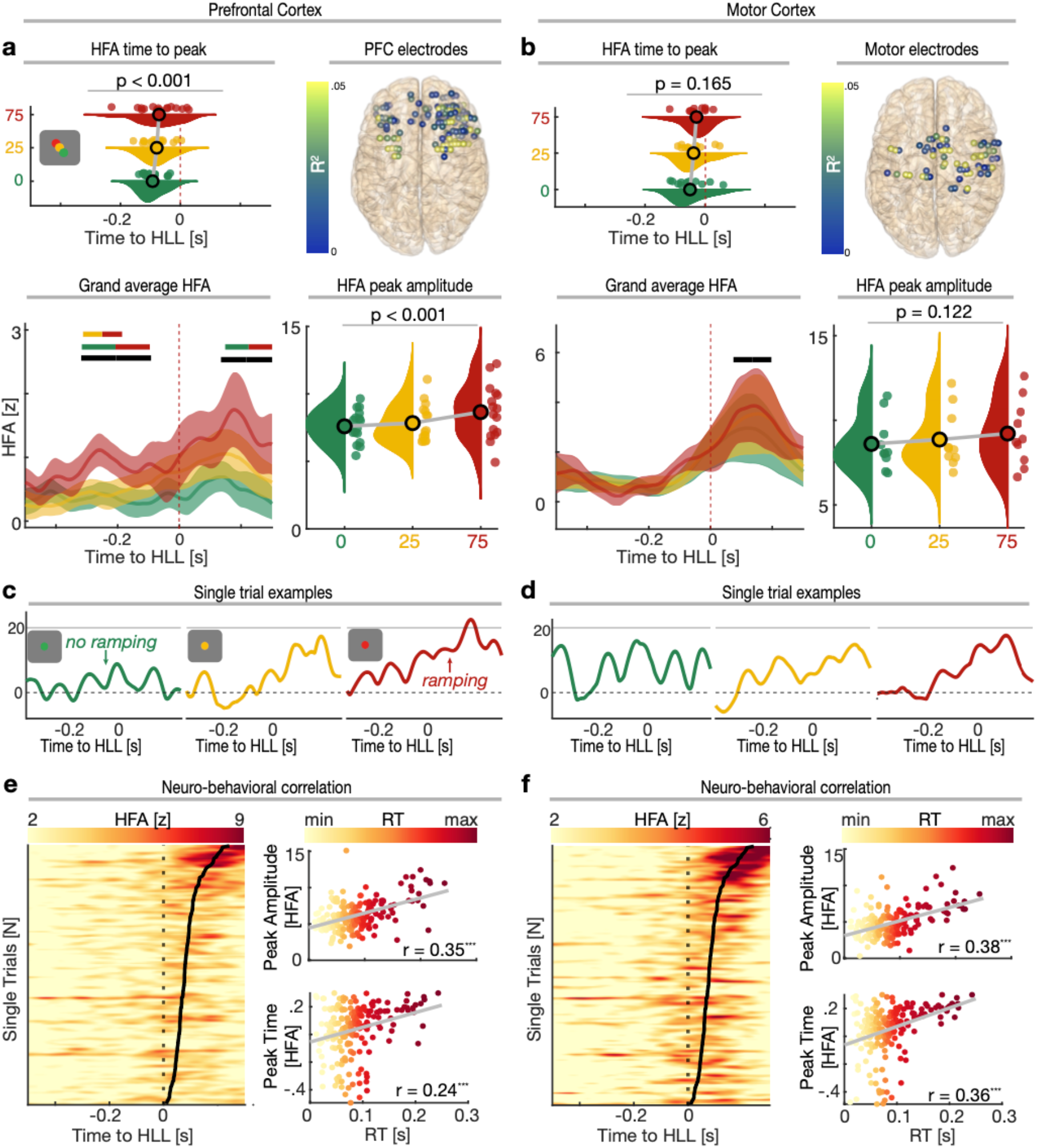
HFA encodes prior evidence and predicts behavior on a trial-by-trial basis. **a**, HFA amplitude (lower right) and peak timing (upper left) both increased with behavioral uncertainty. Lower left: Grand average HFA time courses per context condition (mean ± SEM). Average HFA gradually increased with less predictive context. The single-colored horizontal lines show the temporal extent of significant context-dependent processing. Two-colored horizontal lines indicate the temporal extent of significant clusters obtained from pairwise comparisons. Upper right: Topographical depiction of the neuro-behavioral linear regression. All electrodes are color-coded according to the coefficient of determination (R^2^). **b**, Same as **(a)**, but for motor cortex. Same conventions as in **(a). c-d**, Single trial examples demonstrating the context-dependent change in neural activity. Note the increased ramping dynamics as a function of behavioral uncertainty (from left to right). **e**, Representative single participant example for the neuro-behavioral regression in PFC. Left: Vertically stacked single trials sorted by RT (black line) and color coded according to the z-score. For visualization, panels were smoothed using a 4 trial-wide boxcar function after sorting. Upper right: Relationship between RT and HFA strength. Lower Right: Relationship between RT and HFA peak timing. \*\*\**P* < 0.001. **f**, Same as **(e)**, but for motor cortex. Same conventions as in **(e)**.

We next assessed HFA strength (peak amplitude) and peak timing to quantify neural dynamics on a trial-by-trial basis. We observed that HFA in PFC gradually scaled with behavioral uncertainty (**Fig. 2a**; *F*_2,30_ = 9.77, *P* = 0.0005, 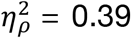; one-way RM-ANOVA). This was not the case for motor cortex (**Fig. 2b**; *F*_2,18_ = 2.37, *P* = 0.122, 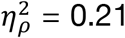). A significant context x ROI interaction confirmed the local specificity of this effect (*F*_2,18_ = 4.67, *P* = 0.046, 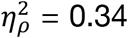; two-way RM-ANOVA). Furthermore, PFC population activity peaked significantly later in trials with high as compared to no uncertainty (**Fig. 2a**; F_2,30_ = 9.07, *P* = 0.0008, 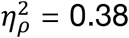; one-way RM-ANOVA). We did not find evidence for such as context-dependent temporal dissociation for motor cortex (*F*_2,20_ = 1.97, *P* = 0.165, 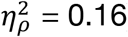). Yet, the direction of the effect was comparable between the two regions (context x ROI interaction; *F*_2,18_ = 1.25, *P* = 0.299, 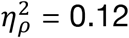; two-way RM-ANOVA). Collectively, these findings indicate that PFC, but not motor cortex, encodes predictive context to guide decisions.

Single-trial associations between HFA amplitude, timing, and behavior were investigated using linear regression. This analysis revealed that HFA in PFC and motor cortex predicted RT on a trial-by-trial basis (**Fig. 2e/f**; *PFC*; *R*^2^ = 0.026, *P* = 6.7 × 10^−18^; *motor cortex*; *R*^2^ = 0.032, *P* = 1.22 × 10^−15^; see **Supplementary Table 1** for partial linear regression). In summary, larger HFA peak amplitudes and slower peak latencies predicted slower RTs. These results highlight delayed and increased HFA responses in states of low predictive context that can be directly mapped to behavior on single trials.

### Ramping dynamics, but not oscillatory signatures dissociate states of uncertainty

We directly tested whether different processing modes implement predictive context (**Fig. 1b**) by disentangling oscillatory and ramping dynamics. We computed the HFA slope on single trials (**Fig. 2c/d**). In line with our main predictions, we found that ramping dynamics were modulated by predictive context. Ramping dynamics in PFC (**Fig. 3a**; *F*_2,30_ = 4.49, *P* = 0.019, 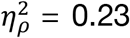; one-way RM-ANOVA), but not in motor cortex (**Fig. 3b**; *F*_2,20_ = 0.36, *P* = 0.698, 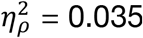), scaled with behavioral uncertainty. Importantly, we did not find support for significant ramping in PFC during trials with no uncertainty (t_15_ = -0.2, *P* = 0.419, Cohen’s *d* = 0.07; one-tailed t-test vs. zero). However, we found significant ramping in PFC during trials with moderate (t_15_ = 3.34, *P* = 0.002, Cohen’s *d* = 1.18) or high uncertainty (t_14_ = 2.15, *P* = 0.024, Cohen’s *d* = 0.79). These results support our prediction that ramping dynamics in PFC are modulated by predictive context.

**Fig. 3.**
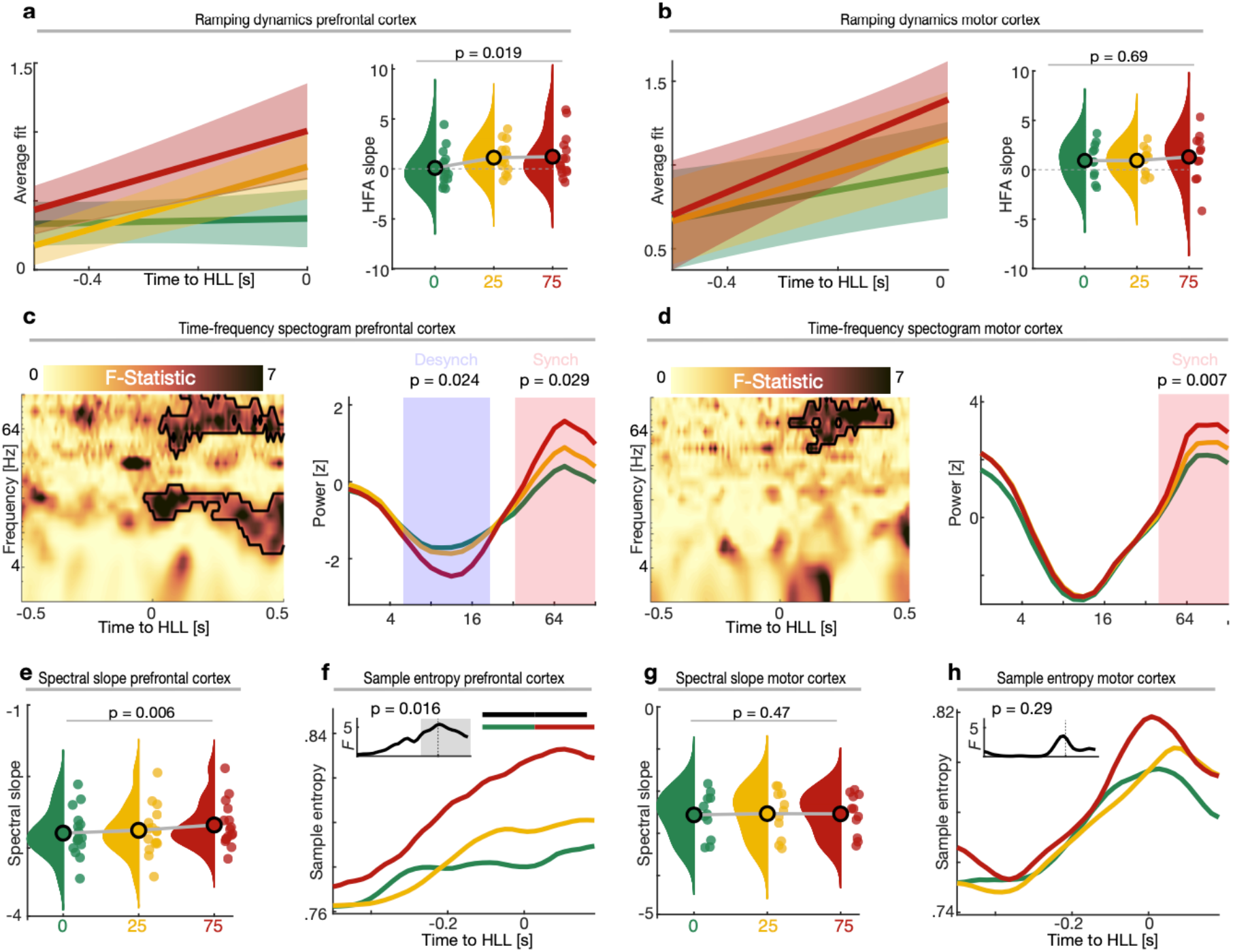
Ramping dynamics dissociate states of behavioral uncertainty and reflect neural excitability. **a**, Left: Grand average linear fit (mean ± SEM) obtained by fitting a linear regression to HFA single trials in PFC. Right: Group-level results depicting the context-dependent modulation of ramping dynamics. **b**, Same as **(a)**, but for motor cortex. Same conventions as in **(a). c**, Time-frequency dynamics in PFC were modulated by predictive context. The black outline indicates the extent of the significant cluster across time and frequency (left panel). Higher frequencies synchronized whereas lower frequencies desynchronized as a function of behavioral uncertainty (right panel). Traces were smoothed for visualization purposes using a 5 Hz running average. **d**, Same as **(c)**, but for motor cortex. Same conventions as in **(c)**. Note the absence of increasing low-frequency desynchronization in motor cortex as a function of behavioral uncertainty. **e**, The aperiodic spectral slope in PFC gets flatter with increasing uncertainty. **f**, Time-resolved PFC sample entropy. The single-colored horizontal lines show the temporal extent of significant context-dependent processing. Two-colored horizontal lines indicate the temporal extent of significant clusters obtained from pairwise comparisons. The small inset depicts the temporal evolution of context-dependent sample entropy. **g**, Same as **(e)**, but for motor cortex. Same conventions as in **(e)**. No context-dependent modulation of the aperiodic slope in motor cortex. **h**, Same as **(f)**, but for motor cortex. Same conventions as in **(f)**. Note the similarly evolving sample entropy across all three predictive context conditions.

Prior studies have argued that ramping dynamics reflect the sequential activation of neural subpopulations with recurrent excitation^25,26^. We therefore examined whether ramping dynamics directly index neural excitability using three surrogate markers of population-level neural excitability (low-frequency desynchronization, spectral exponent, and sample entropy^27-30^). While high-frequency synchronization was evident in both PFC (**Fig. 3c**; *F*_2,30_ = 849.3, *P* = 0.029; cluster test) and motor cortex (**Fig. 3d**; *F*_2,20_ = 743.3, *P* = 0.007), low-frequency desynchronization was only apparent in PFC (**Fig. 3c**; *F*_2,30_ = 922.1, *P* = 0.024). A context x ROI interaction effect in the lower frequencies (2-19 Hz; t_10_ = -548.72, *P* = 0.006) confirmed that the low-frequency desynchronization during states of high uncertainty was specific to PFC. The spectral slope has been shown to closely track the excitation/inhibition (E/I) balance in neural circuits (flatter slopes indicate more excitation^29,31^). We found that the spectral slope got flatter with behavioral uncertainty in PFC (**Fig. 3e**; *F*_2,30_ = 5.97, *P* = 0.006, 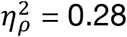; one-way RM-ANOVA), but not motor cortex (**Fig. 3g**; *F*_2,20_ = 0.77, *P* = 0.476, 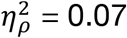; context x ROI interaction effect; *F*_2,20_ = 14.2, *P* = 0.0002, 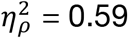; two-way RM-ANOVA). We computed time-resolved fluctuations in E/I balance using sample entropy^30^. Time-resolved sample entropy was context-dependent in PFC and showed the strongest increase in trials with high uncertainty (**Fig. 3f**; *F*_2,30_ = 68.32, *P* = 0.016; cluster-test). In contrast, we found no evidence of time-resolved entropy in motor cortex to be context-dependent (**Fig. 3h**; *F*_2,20_ = 4.01, *P* = 0.291). No context x ROI interaction was present (no cluster at p < 0.05). Collectively, this set of findings demonstrates that predictive context initiates a shift in ramping dynamics and neural excitability during evidence accumulation. We show that these shifts are most pronounced in PFC (**Fig. 3a/c/e/f**) in comparison to motor cortex (**Fig. 3b/d/g/h**) and gradually scale with behavioral uncertainty.

Next, we investigated how oscillatory dynamics were modulated by predictive context during evidence accumulation. We extracted all HFA peaks (**Fig. 4a**) and performed peak-triggered averaging (PTA; **Fig. 4b**). We found that HFA is nested in a theta oscillation (~5 Hz; **Fig. 4b**). In order to quantify this on a group level and assess context-dependent modulations, we spectrally decomposed the PTA and separated oscillatory from aperiodic background activity by means of irregular-resampling auto-spectral analysis (IRASA)^32^. A cluster-based permutation test revealed reduced oscillatory power in PFC during trials with high uncertainty (**Fig. 4c;** first cluster; *F*_2,30_ = 58.16, *P* = 0.0009; second cluster; *F*_2,30_ = 56.54, *P* = 0.0009; cluster test). Importantly, this context-dependent modulation was not driven by changes in the peak frequency of the theta oscillations (**Fig. 4c**). Pronounced theta peaks were present irrespective of the contextual cue. We also inferred the instantaneous peak frequency directly on the HFA signal by computing the interval between adjacent HFA peaks (**Fig. 4d**)^33^. The instantaneous HFA peak frequency decreased with uncertainty (**Fig. 4e/f**; *F*_2,30_ = 4.14, *P* = 0.025, 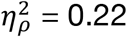; one-way RM-ANOVA). While theta oscillatory peaks were equally present in motor cortex, we found no context-dependent modulation in either the oscillatory power of the PTA (**Supplementary Fig. 2**; *F*_2,20_ = 7.61, *P* = 0.368; cluster test) or the instantaneous frequency (**Supplementary Fig. 2**; *F*_2,20_ = 2.24, *P* = 0.132, 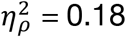; one-way RM-ANOVA) of the HFA signal. Taken together, we did not find strong evidence for a context-dependent modulation of oscillatory power. In contrast to the presumed switch from an oscillatory to a continuous processing regime (**Fig. 1b**), we found that neural oscillations are ubiquitous across all predictive contexts.

**Fig. 4.**
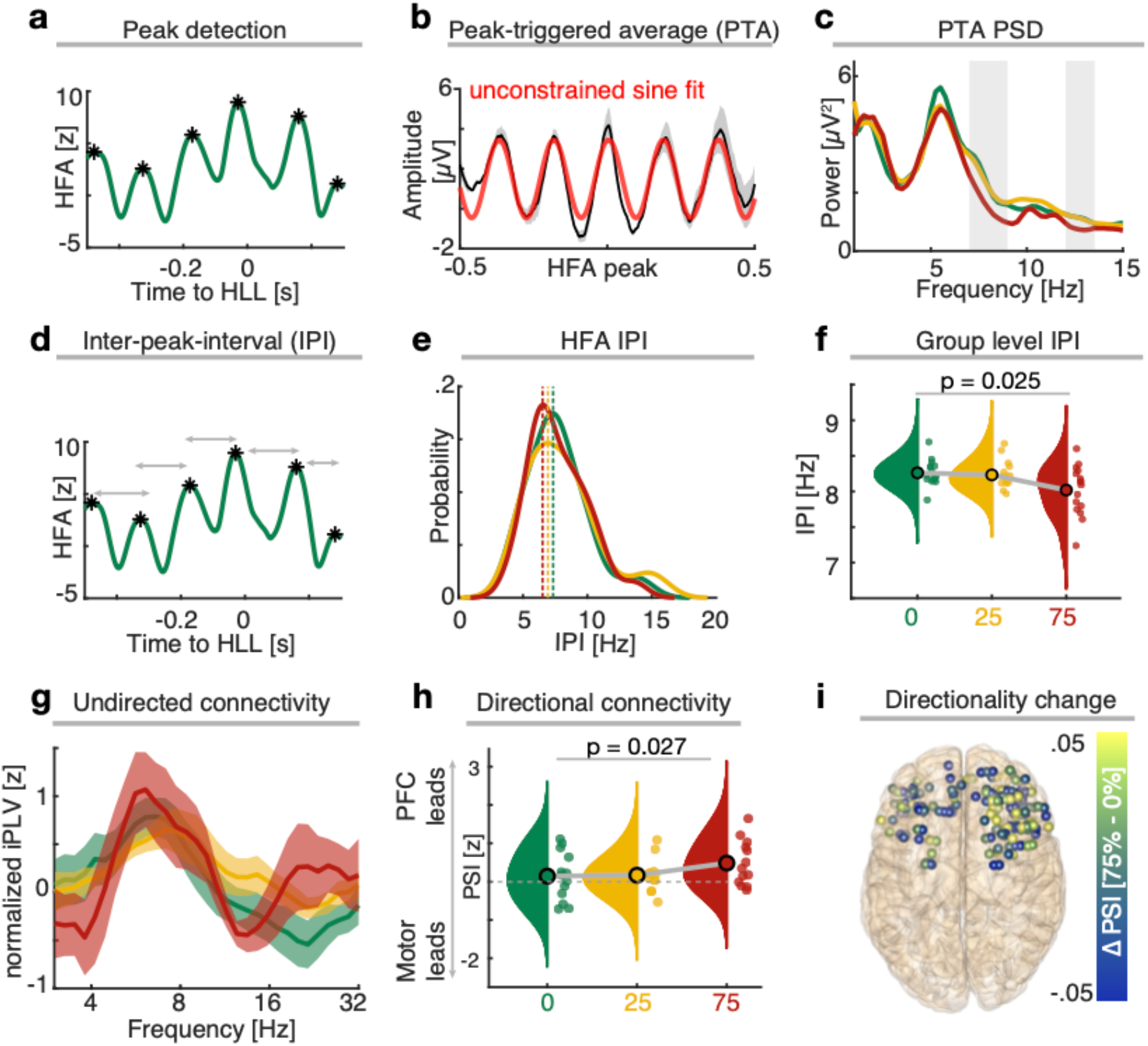
Theta oscillations modulate HFA and mediate context-dependent information flow from PFC to motor cortex. **a**, Example of the peak detection on single trial HFA traces (black asterisk). Note the waxing and waning pattern in single trials. **b**, Peak-triggered average (PTA; mean ± SEM; ±0.5s from HFA peak) in a representative single participant across PFC electrodes. HFA was nested into a ~5 Hz theta oscillation (red line depicts a sine fit to the PTA). **c**, Grand average 1/f-corrected power spectrum computed on the PTA time-series using IRASA. Shaded grey areas depict the extent of significant context-dependent power modulation. Pronounced theta peaks were present in all predictive context conditions. **d**, Example trace depicting the quantification of the inter-peak interval (IPI) as a time length between two contiguous peaks. **e**, Single electrode example showing the IPI distribution across conditions. Vertical dashed lines represent the peaks of the distributions. **f**, Reduced IPI with increasing behavioral uncertainty. **g**, Grand-average (mean ± SEM) prefrontal-motor undirected connectivity. Undirected connectivity was not modulated by states of uncertainty, but showed pronounced peak connectivity in the theta band. **h**, Directional prefrontal-motor connectivity in the theta band. Directional information flow from PFC to motor cortex was enhanced during states of high uncertainty. **i**, Topographical depiction of the directional change in information flow from PFC to motor cortex between distinct states of uncertainty.

This set of findings raised the question which role neural oscillations play in processes where evidence needs to be converted into an action. Based on the well-established role of neural oscillations in mediating inter-areal communication^6,34^, we tested whether oscillations synchronize the prefrontal-motor network. We computed the imaginary phase-locking value (iPLV) between prefrontal-motor electrode pairs to assess network connectivity. We observed strong prefrontal-motor synchrony in the theta band (**Fig. 4g**; 6.4 ± 1.3 Hz; mean ± SD) with no difference between context conditions (*F*_2,26_ = 3.82, *P* = 0.39; cluster test). To assess directional interactions, we computed the phase-slope index (PSI)^35^. We first identified the individual iPLV peak frequency for every prefrontal-motor electrode pair prior to computation of the PSI. We found that directional theta connectivity from PFC to motor cortex was context-dependent (**Fig. 4h/i**; *F*_2,24_ = 4.2, *P* = 0.027, 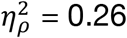; one-way RM-ANOVA) and strongest in trials with high behavioral uncertainty (t_24_ = 2.95, *P* = 0.006, Cohen’s *d* = 1.16; two-tailed t-test vs. zero). Collectively, this set of findings demonstrates that ramping dynamics in PFC dissociate states of behavioral uncertainty while neural oscillations dynamically coordinate the prefrontal-motor network interaction in a context-dependent manner.

### Population dynamics in PFC encode context and actions in orthogonal subspaces

While it is well established that neural oscillations reflect coordinated population activity^36^, the population correlates of ramping dynamics are understood to a lesser degree. Having established that ramping dynamics likely reflect a mechanism of context-dependent evidence accumulation, we assessed how ramping dynamics impact population dynamics. In neural state space, population dynamics can be conceptualized as a trajectory across time through a *N*-dimensional space (Methods). We computed the multidimensional distance (MDD) between pairwise trajectories. We observed that neural state-space trajectories in PFC were context-dependent and diverged prior to participants’ choice of behavioral response (button release) (**Fig. 5a**; t_16_ = 46.13, *P* = 0.024; cluster test). This effect was driven by strong state transitions in trials with high behavioral uncertainty (**Fig. 5a**; *F*_2,32_ = 96.01, *P* = 0.011). No evidence was found for a context-dependent evolution of neural trajectories in motor cortex (**Fig. 5b**; t_13_ = 13.69, *P* = 0.116). Instead, the neural state transitions revealed a highly similar pattern across all three levels of uncertainty (**Fig. 5b**; *F*_2,26_ = 23.81, *P* = 0.123).

**Fig. 5.**
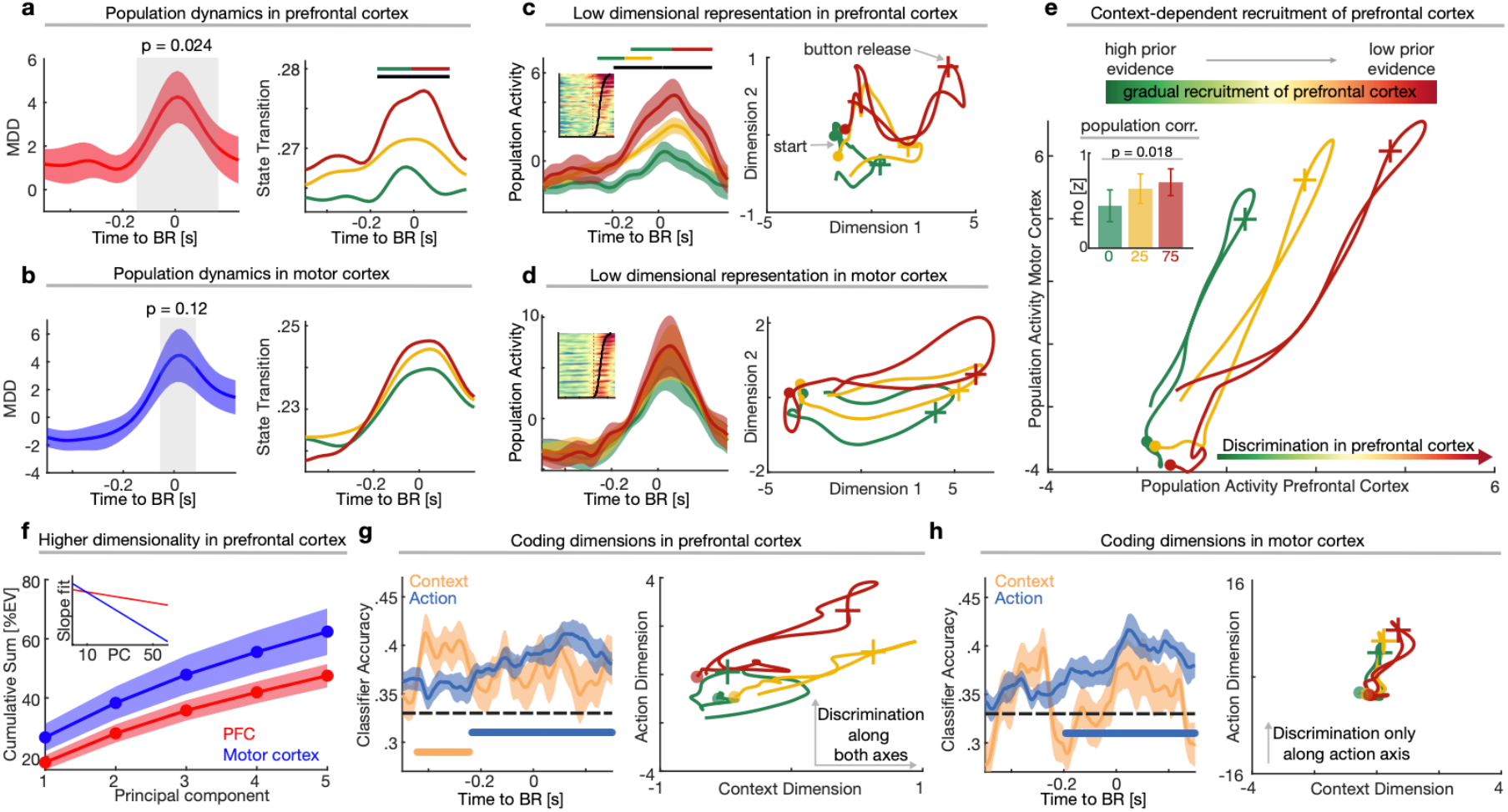
Low-dimensional representation of context and action in orthogonal coding dimensions. **a**, Left: Normalized multidimensional distance (MDD; mean ± SEM; Methods) between the three types of predictive context in the neural state space (PFC). Significant divergence of neural trajectories prior to the behavioral response. Shaded grey areas depict the temporal extent of significant clusters. Right: Within-condition transitions in activation states. Note the strong vs. weak neural state transitions in trials with high vs. no behavioral uncertainty, respectively. The single-colored horizontal lines show the temporal extent of a significant context-dependent dissociation. Two-colored horizontal lines indicate the temporal extent of significant clusters obtained from pairwise comparisons. **b**, Left: Normalized MDD (mean ± SEM) between the three context conditions within motor neural state space. Neural trajectories were statistically indistinguishable. Right: The dynamics of neural activation state transitions revealed a similar profile across all context conditions with strong state changes until the participants’ response. **c**, Left: Context conditions can be dissociated based on the first principal component (PC1) of the HFA, reflecting the most dominant pattern of population activity. Horizontal lines share the same convention as in **(a)**. Population activity gradually increased with uncertainty. The inset depicts stacked single trials of the population activity sorted by RT (black line). Population activity explained a significant proportion of behavioral variance on single trials (linear regression; *R*^2^ = 0.02, *P* = 7.9 × 10^−15^). Right: Low-dimensional neural trajectories formed by the top two PCs in PFC. The filled dots indicate the start (−0.5s prior to BR) and the crosses indicate the moment of BR. Note the joint activation pattern at the start of the trial, but the progressively increasing divergence of context-dependent neural trajectories until the participants’ response. States of high behavioral uncertainty show a more dynamic profile with a more complex pattern through the neural state space. Neural trajectories were smoothed using a 50 ms running average. **d**, Left: Context conditions were not dissociable based on the first principal component (PC1) of the HFA, reflecting the most dominant pattern of population activity in motor cortex. Note the highly similar activation profiles across all three predictive context conditions (as compared to the context-dependent dynamics in PFC). The inset depicts stacked single trials of the population activity sorted by RT (black line). Population activity still captured a significant proportion of behavioral variance on single trials (linear regression; *R*^2^ = 0.01, *P* = 1.9 × 10^−7^). Right: Low-dimensional neural trajectories formed by the top two PCs in motor cortex. Same convention as in right panel of **(b)**. All three neural trajectories follow a similar curved path through state space. **e**, Projection of the population activity in PFC and motor cortex into a common two-dimensional space. Each point reflects the joint activation state of population activity in the prefrontal-motor network. Note that the main discrimination of the three types of predictive context occurs in PFC, suggesting a gradual recruitment of PFC activity with increasing uncertainty. Axes are equally scaled for comparison. Filled dots and crosses, same conventions as in **(b)** and **(d)**. The inset displays the z-transformed (Methods) power correlation coefficient between prefrontal-motor population activity per context condition. Functional connectivity between the population activity in the prefrontal-motor network increased with uncertainty. **f**, Cumulative percent variance (%EV; mean ± SEM) explained by the top five PCs in PFC (red) and motor cortex (blue). The cumulative %EV is significantly lower in PFC. Similarly, the variance explained per PC decayed more rapidly in motor cortex (inset), highlighting increased dimensionality in PFC. **g**, Left: Grand average decoding accuracy (mean ± SEM) for context and action within PFC (Methods). Horizontal lines indicate the extent of significant temporal clusters (color coded for the respective feature). Maximal context classification emerged prior to action classification. Right: Joint representation of the neural trajectories formed the two coding dimensions (context and action; Methods). Each point represents the joint activation state at time *t* formed by the two coding dimensions. Note that the neural trajectory does not progress along the context dimension in trials with no uncertainty, suggesting a direct transformation of predictive context into an action plan at trial start. **h**, Left: Grand average decoding accuracy (mean ± SEM) for context and action within motor cortex. Only action, but not context, could be decoded in motor cortex. Same conventions as in **(g)**. Right: Joint representation of the neural trajectories formed the two coding dimensions. Note that all neural trajectories progress along the action, but not the context dimension. Axes in left panels of **(g)** and **(h)** are equally scaled for comparison (y-axis = four times x-axis).

We extracted the most dominant, population-wide activity pattern using principal component analysis (PCA) to examine the latent dynamics underlying context-dependent evidence accumulation (**Fig. 5c/d/e**). We found that the top three principal components (PCs) captured 35.76 ± 14.15% of the variance (mean ± SD) in PFC and 50.6 ± 24.88% of the variance in motor cortex (**Fig. 5f**; t_13_ = - 2.2, *P* = 0.046, Cohen’s *d* = 0.66; two-tailed t-test). We also observed that the variance explained per PC decreased more rapidly in motor cortex as compared to PFC (t_13_ = -2.01, *P* = 0.065, Cohen’s *d* = 0.67), suggesting that neural dynamics are higher dimensional in the human PFC than motor cortex. In line with this finding, population activity in PFC (PC1) revealed context-dependent dynamics (**Fig. 5c**; *F*_2,32_ = 1.18 × 10^3^, *P* = 0.008; cluster test) with larger activation states in trials with high behavioral uncertainty. In comparison, we found similar latent dynamics across all context conditions in motor cortex (**Fig. 5d**; *F*_2,26_ = 230.85, *P* = 0.079). This indicates that PFC, but not motor cortex, is gradually recruited as a function of behavioral uncertainty (**Fig. 5e**). Next, we tested whether predictive context also modulates the inter-areal coupling in the prefrontal-motor network on a population level. We therefore quantified the degree of functional connectivity between population activity in PFC and motor cortex on a single-trial level (Methods). This revealed that the strength of the prefrontal-motor interaction scaled with behavioral uncertainty (inset **Fig. 5e**; *F*_2,26_ = 4.69, *P* = 0.018, 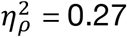; one-way RM-ANOVA) and explained behavior on a single trial basis (*R*^2^ = 0.014, *P* = 2.4 × 10^−9^; linear regression).

We specifically assessed which latent dimension reflects contextual encoding in human PFC, given that neural dynamics were high-dimensional in PFC. To determine the relevant coding dimensions, we employed multivariate pattern classifiers (linear discriminant analysis; LDA) in PC space separately for both regions. This approach defined the coding dimensions that maximally discriminated context and behavioral performance (RT; split into terciles; Methods), thus, dissociating dynamics that mediated contextual processing and subsequent action execution. We found that human PFC encoded both context and action (**Fig. 5g**; context: t_16_ = 352.99, *P* = 5.9 × 10^−4^, Cohen’s *d* = 0.83; action: t_13_ = 1445.56, *P* = 1.9 × 10^−4^, Cohen’s *d* = 1.4; cluster tests). Critically, context could be decoded prior to action (**Fig. 5g**; context: 443ms prior to button release; action: 234ms prior to button release), suggesting that context was integrated before being translated into an action plan. Importantly, we found orthogonal coding dimensions for context and action coding (maximal discrimination in distinct PCs) in 11/14 participants (*P* = 0.057, Binomial test). In contrast, we were only able to reliably discriminate action, but not context, from motor cortex (**Fig. 5h**; context: t_13_ = 39.32, *P* = 0.485, Cohen’s *d* = 0.75; action: t_9_ = 1118.44, *P* = 1.9 × 10^−4^, Cohen’s *d* = 1.39), which indicates that the relevant contextual computations were completed at the level of the output stage.

Collectively, we have shown that the activation state of population dynamics in the human PFC gradually scales as a function of behavioral uncertainty. In a final analysis step, we characterized how population dynamics interact with neural oscillations to support goal-directed behavior. We therefore extracted the dimension with the strongest oscillatory theta power in every participant (**Fig. 6a/b**). Next, we employed LDA classifiers to assess the coding features of the theta component. In both PFC and motor cortex, we found that the dimension with the strongest theta power significantly coded action (**Fig. 6c**; PFC; t_13_ = 740.72, *P* = 0.0004, Cohen’s *d* = 0.82; motor cortex; t_9_ = 816.03, *P* = 0.002, Cohen’s *d* = 1.32; cluster tests), but not context (PFC and motor cortex; no cluster at p < 0.05). In support of this observation, we found no evidence that the theta dimension and the previously determined action dimensions are embedded in distinct subspaces (*P* = 0.18, Binomial test). Remarkably, the dimension with the strongest theta power most likely matched PC1 (PFC: 11/17 participants; motor cortex: 11/14 participants). Finally, we observed that neural dynamics embedded in the dimensions with strongest theta power are functionally coupled within the prefrontal-motor network (**Fig. 6d**; *P* = 0.003, Cohen’s *d* = 0.69; Wilcoxon rank sum test). However, coupling strength was not modulated by predictive context (*F*_2,26_ = 1.36, *P* = 0.273, 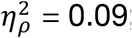; one-way RM-ANOVA), indicating a functional role of theta oscillations to mediate the hand-off of action plans from prefrontal to motor cortex. Taken together, these findings reveal that structured population activity in PFC encodes and integrates predictive context into a higher-level action plan that is executed at the level of motor cortex. Our results demonstrate that the transformation from PFC-dependent context integration to goal-directed action execution in motor cortex is mediated by directed theta-band connectivity (cf. **Fig. 4g-i**).

**Fig. 6.**
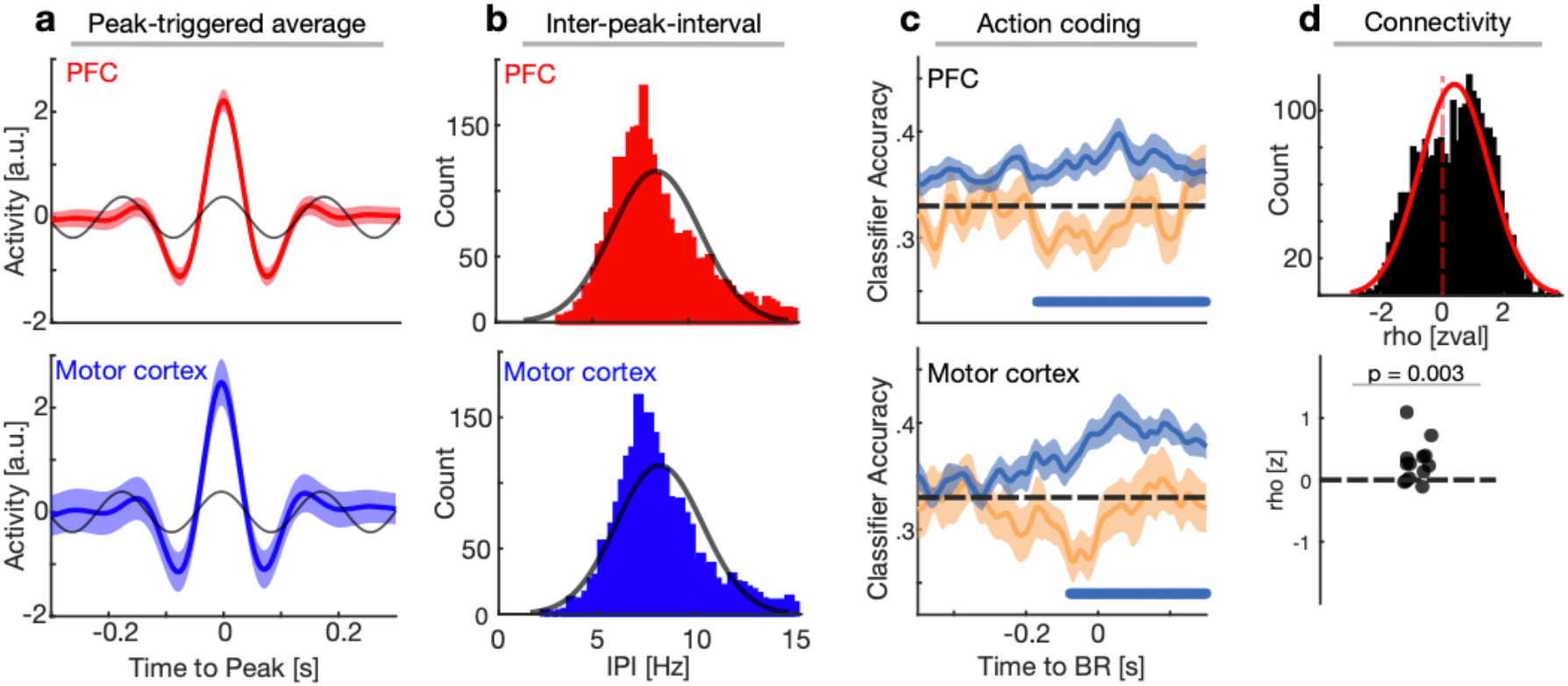
Theta oscillations temporally coordinate action-encoding subspaces in the prefrontal-motor network. **a**, The peak-triggered average (PTA) across participants obtained from the PC with the strongest theta power in PFC (upper panel) and motor cortex (lower panel). The black lines depict a sine fit to the PTAs. **b**, The instantaneous frequency of the identified theta PC as computed by the inter-peak-interval (upper panel shows the distribution for PFC, the lower panel for motor cortex). Both PFC and motor cortex showed strong oscillatory peaks in the theta-frequency band. **c**, Grand average decoding accuracy (mean ± SEM) for context and action within the identified theta PC in PFC (upper panel) and motor cortex (lower panel). The theta dimension significantly coded action, but not context in both PFC and motor cortex. **d**, Upper panel: Histogram depicting z-normalized power correlation coefficients between the theta PCs in PFC and motor cortex. Lower panel: Significant coupling between the theta PCs in PFC and motor cortex. Single dots represent the z-normalized (permutation) power correlation coefficients.

## Discussion

Context-guided decision-making is a hallmark of flexible human behavior. To date, it remains unknown how contextual priors are encoded to guide decision processes in humans. Previous work in NHP indicated that (pre-)motor cortex might mediate context-dependent decision-making^10,12-15^. While earlier theories, such as the active sensing framework^4,5^, emphasized that neural coding is mainly reflected in local activity profiles (i.e. neural firing or oscillatory (de-)synchronization), novel population-based theories now suggest that context-dependent processing is distributed across large-scale neural populations^10,12^.

Thus far, the population doctrine had its greatest impact on understanding movement-related computations in the motor system^37-40^, but it might also provide a powerful framework to understand higher cognitive processes^41^. Using a predictive motor task, we demonstrate that (I) behavioral uncertainty is reflected in neural indices of uncertainty as quantified by uni- (**Fig. 2/3**) and multivariate analyses (**Fig. 5**). In line with the active sensing framework, we show that (II) behavioral uncertainty introduces a shift from an *oscillatory* to a continuous *ramping* processing mode (**Fig. 3/4**). Using population-based analysis strategies, we demonstrate that (III) oscillatory and ramping dynamics reflect dissociable population signatures that support distinct aspects of encoding, transmission and execution of context-dependent action plans (**Fig. 5**). Specifically, we show that (IV) prefrontal population activity encodes predictive context and action plans in serially unfolding orthogonal subspaces, while we observed motor cortex to encode action plans only (**Fig. 5**). Furthermore, our results reveal that (V) theta synchrony temporally coordinates action-encoding population subspaces, thereby mediating the hand-off of action plans from prefrontal to motor cortex (**Fig. 6**). Collectively, our results demonstrate how two hallmarks of large-scale population activity, namely continuous population dynamics and oscillatory synchrony, operate in concert to guide context-dependent human behavior.

### Oscillatory and ramping dynamics reflect distinct population signatures of context-dependent behavior

The influential active sensing framework postulates that the brain switches from an energy-efficient *oscillatory* processing mode during states of high predictability to an energy-consuming continuous *ramping* processing mode in states of low predictability. Evidence for this theory has mainly been obtained in NHP auditory cortex^7,8^, but it had been argued that similar principles apply to higher-order cortical areas^5^. In line with this framework, we found that the transition from high to low prior evidence increased ramping dynamics in the human PFC, but not in motor cortex. Contrary to the theory, we did not find evidence for a modulation of local oscillatory dynamics as a function of predictability. In addition, a related line of inquiry argued that frontal theta activity constitutes a mechanism of cognitive control, especially in states of high uncertainty^22,42^. Using direct brain recordings in humans, we found that directional theta synchrony is inversely related to predictability. We found stronger directional theta synchrony from prefrontal to motor cortex in states of high uncertainty, indicating a flexible recruitment and network engagement when limited predictive context is available. Using population-based decoding, we found that theta oscillations were not associated with the encoding of predictions per se, but that theta activity was confined to the action subspaces of the population activity. This finding is in line with the communication-trough-coherence hypothesis^6^. Moreover, it indicates that the PFC encodes task-relevant context, devises the appropriate motor plan, and hands off the action plan to the motor cortex for execution through theta synchrony. Collectively, these results demonstrate that several hallmarks of predictive processing that have primarily been captured using univariate metrics in fact reflect coordinated and functionally specialized, population-wide activity patterns.

Translating findings that were obtained in NHP into human research is hampered by the fact that signals from different recording modalities are typically being compared (e.g. single unit vs. EEG activity). Here, we analyzed HFA in humans as a proxy of multiunit activity firing^17-19^. HFA offers the advantage that it already constitutes an aggregate metric that summarizes the underlying population activity. Recent work demonstrated that HFA contains more behaviorally relevant information than single-/multi-unit activity or EEG activity, and therefore constitutes a highly suitable level of abstraction to study population-wide activity^43^. Furthermore, theta oscillations temporally structure HFA activity through phase-amplitude cross-frequency coupling, and thus resemble previous findings that neural firing is controlled by network oscillations^20,44^.

### The population doctrine and cognitive processing

The population doctrine is an emerging concept highlighting that population activity, and not the single unit per se, reflects the essential unit of computation in the brain^16,45^. Population activity has mainly been studied in NHP (pre-)motor cortex where distinct movement trajectories are represented by unique neural trajectories of the population^37,40^. While previous evidence in NHP indicated that (pre-)motor cortex performs context-dependent computations^13-15,46^, we found that neural trajectories in prefrontal, but not motor cortex, dissociated the current predictive context. Critically, we observed that large-magnitude neural states within PFC indexed behavioral uncertainty. We found that PFC settled into a low-energy state (smaller magnitude, only covering a limited subspace of the entire state space) during states of high predictability. Critically, these patterns could only be observed when using multivariate analysis strategies (**Fig. 5**; c.f. **Fig 2** for the univariate approach) that take coordinated variability across different recording sites into account.

Previous work in NHP demonstrated that motor cortex exhibits a low-dimensional structure^45,47^. Here, we replicate this finding in humans, but in contrast to NHPs, we found no evidence that the human motor cortex encodes predictive context. We observed that motor cortex relies on input from PFC, which encodes both context as well as the current action plan. Importantly, prefrontal population activity is high-dimensional in nature, where distinct operations (context encoding and action planning) are encoded in orthogonal subspaces. We argue that the high-dimensional prefrontal functional architecture constitutes a substrate for flexible goal-directed behavior and that simultaneous processing in separate coding dimensions maximizes information-coding capacity of the underlying population.

## Conclusions

In the present study, we demonstrate that high-dimensional prefrontal population dynamics encode predictive context and action plans in orthogonal subspaces. We demonstrate that a lack of prior evidence comes at a behavioral (increased response time/error rates) as well as a neural (large magnitude neural states) cost. Moreover, our results reveal a functional dissociation of population trajectories and oscillatory synchrony, and indicate a division of labor between prefrontal and motor cortex. Specifically, we found that prefrontal population trajectories encode behaviorally relevant variables, while oscillatory synchrony mediates the prefrontal-motor transmission of action plans. We observed low-dimensional neural dynamics in human motor cortex, which did not encode predictive context, but relied on theta-mediated input from higher-order prefrontal areas. We studied context-dependent motor behavior using univariate as well as multivariate analyses and thereby demonstrated that ramping and oscillatory signatures of predictive processing in fact constitute dissociable signatures of coordinated population activity underlying flexible human behavior. These findings pave the way for future studies to understand human goal-directed behavior and provide the first demonstration that population dynamics and oscillatory synchrony interact in concert to guide flexible human behavior.

## Online Methods

### Patients and implantation procedure

We obtained intracranial recordings from a total of 19 pharmaco-resistant epilepsy patients (33.73 years ± 12.52, mean ± SD; 7 females) who underwent presurgical monitoring and were implanted with intracranial depth electrodes (DIXI Medical, France). Data from one patient were excluded from neural analyses because a low-pass filter was applied at 50 Hz during data export from the clinical system, thus, precluding analyses focusing on HFA. All patients were recruited from the Department of Neurosurgery, Oslo University Hospital. Electrode implantation site was solely determined by clinical considerations and all patients provided written informed consent to participate in the study. All procedures were approved by the Regional Committees for Medical and Health Research Ethics, Region North Norway (#2015/175) and the Data Protection Officer at the Oslo University Hospital as well as the University Medical Center Tuebingen (049/2020BO2) and conducted in accordance with the Declaration of Helsinki.

### iEEG data acquisition

Intracranial EEG data were acquired at the Oslo University Hospital at a sampling frequency of 512 Hz using the NicoletOne (Nicolet, Natus Neurology Inc., USA) or at a sampling frequency of 16 KHz using the ATLAS (Neuralynx) recording system.

### CT and MRI data acquisition

We obtained anonymized postoperative CT scans and pre-surgical MRI scans, which were routinely acquired during clinical care.

### Electrode localization

Two independent neurologists visually determined all electrode positions based on individual scans in native space. For further visualization, we reconstructed the electrode positions as outlined recently^48^. In brief, the pre-implant MRI and the post-implant CT were transformed into Talairach space. Then we segmented the MRI using Freesurfer 5.3.0^49^ and co-registered the T1 to the CT. 3D electrode coordinates were determined using the Fieldtrip toolbox^50^ on the CT scan. Then we warped the aligned electrodes onto a template brain in MNI space for group-level analyses.

### Task

Participants performed a predictive motor task where they had to continuously track a moving target and respond as soon as the target hits or withhold their response if the target stops prior a predefined spatial position using their dominant hand (**Fig 1a**). Prior to the main experiment, participants were familiarized with the task by means of a short practice session. Each trial started with a baseline period of 500ms followed by a cue (presented for 800ms centered) that informed participants about the likelihood that the moving target would stop prior to the lower limit (hit lower limit; HLL; **Fig. 1a**). Thus, the predictive cue could be directly translated into the probability that either of two possible action scenarios will occur: button release (BR) vs. withhold response (Bernoulli distribution). Participants were instructed to either release the button as soon as the target hits (“Go” trials) or withhold their response if the target stops prior to the HLL (“Stop” trials). We parametrically modulated the likelihood of stopping. A green circle indicated a 0% likelihood, an orange circle indicated a 25% likelihood and a red circle indicated a 75% likelihood that the moving target would stop prior to the HLL. Hence, participants were able to fully predict the outcome on trials with a 0% likelihood and already prepare the motor response. However, in trials with a 25% or 75% likelihood of stopping, they continuously had to accumulate evidence in order to decide whether to release the button or withhold the response. Upon receiving the predictive cue, participants were able to start the trial in a self-paced manner by pressing the space bar on the keyboard. By pressing the space bar, the target would start moving upwards and reach the HLL after 560 – 580ms. The upper boundary was reached after 740 – 760ms, thus, leaving 160ms between the HLL and the upper boundary. If participants released the button within this 160ms interval, the trial was considered as correct. Trials in which the button was released either before or after this interval were considered as incorrect. Feedback on trial performance was provided upon each trial for 1000ms.

### Behavioral data analysis

We quantified reaction time (RT) as the time passed between the moving target reaching the HLL and the participants’ response. We considered both correct and incorrect trials in our analyses on RT. Accuracy was quantified as the average number of correct responses relative to the number of trials. We used the interquartile range (IQR) as a measure for behavioral trial-by-trial variability^51^. We also considered the signal detection theoretic measures *d’* (d-prime) and *c* (criterion)^52^. While *d’* quantifies the distance between the signal (e.g. go trials) and noise distribution (e.g. stop trials), *c* reflects a participant’s propensity to choose yes or no (decision criterion). Due to the nature of the task (absence of noise distribution in the 0% condition), we were only able to quantify *d’* and *c* for conditions with a 25% or 75% likelihood of stopping.

### Intracranial EEG analysis

#### Preprocessing and artifact rejection

Intracranial EEG data were demeaned, linearly de-trended, locally re-referenced (bipolar derivations to the next adjacent lateral contact) and if necessary down-sampled to 512 Hz. To remove line noise, data were notch-filtered at 50 Hz and all harmonics. Subsequently, a neurologist visually inspected the raw data for epileptic activity. Channels or epochs with interictal epileptic discharges (IEDs) and other artifacts were removed.

#### Trial definition

We extracted 10 seconds long, partially overlapping trials to prevent edge artifacts in subsequent filtering. We excluded all stop trials and focused subsequent analyses on go trials. Trials were event-locked to the HLL unless otherwise stated.

#### Definition of regions of interest

The pre-selection of electrodes was guided by our question on how the human prefrontal-motor network is engaged during context-dependent evidence accumulation. Electrodes were classified into discrete PFC and motor ROIs based on surface anatomy using the Anatomical Automatic Labeling atlas (ROI_MNI_V4.nii)^53^. Electrodes in the following areas were considered to be in the PFC ROI (equal for both hemispheres): superior frontal gyrus (orbital, medial and dorsolateral part), medial frontal gyrus, inferior frontal gyrus (opercular, triangular and orbital part). Electrodes in the following areas were considered to be motor electrodes (equal for both hemispheres): precentral gyrus, supplementary motor area, paracentral lobule. In total, 17 patients were implanted with clean, artifact-free electrodes in PFC, 14 patients in motor cortex and 14 patients were implanted with clean, artifact-free electrodes in both ROIs.

#### HFA extraction

The extraction of the high-frequency activity time series was conducted in a three-step process. In the first step, we bandpass-filtered the raw data epochs (10 seconds) between 70-150 Hz into eight, non-overlapping 10 Hz wide bins. We then applied the Hilbert transform to obtain the instantaneous amplitude of the filtered time series. In a last step, we normalized the high-gamma traces using a bootstrapped baseline distribution^54,55^. This involved randomly resampling baseline values (from -0.2 to -0.01s relative to cue onset) 1000 times with replacement and normalizing single high-gamma traces by subtracting the mean and dividing by the standard deviation of the bootstrap distribution. The high-gamma traces were finally averaged across the eight bins. This procedure mitigates the effect of the 1/f power drop-off and enables comparable estimates across different conditions by minimizing the influence of different baseline distributions onto task-related activity.

#### Context-dependent neural information

We identified context-encoding electrodes using a well-established information theoretical approach that has been used in both human and non-human primate studies^22-24,56^. We employed a one-way analysis of variance (ANOVA) to quantify the percentage of HFA variance that could be explained by our behavioral regressor “predictive context”. The amount of percent explained variance was quantified using ω^2^ as

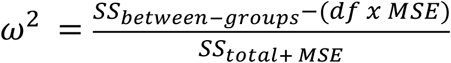

where *SS*_*total*_ reflects the total sum of squares across *n* trials,

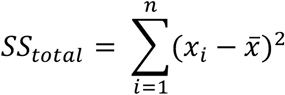

*SS*_*between−groups*_ the sum of squares between *G* groups (e.g. factor levels),

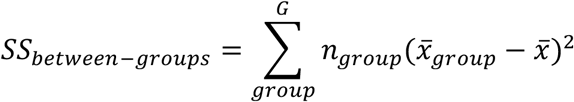

*MSE* the mean square error,

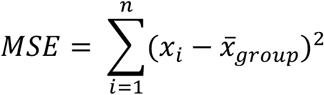

and *df* the degrees of freedom specified as *df = G* − 1. In order to obtain a time series of context-dependent neural information, we estimated ω^2^ using a sliding window of 50ms that was shifted in steps of 2ms. Electrodes that exhibited a significant main effect of predictive context for at least 10% of the trial length were defined as context-encoding electrodes. Note that this approach was blind with respect to both direction and timing of the effect. Finally, to minimize inter-individual variance and maximize the sensitivity to identify a temporally consistent pattern that accounts for most of the variance explained by predictive context within the context-encoding electrodes across participants, we used principal component analysis (PCA)^24,57^. PCA was applied to the *F* value time series concatenated across participants (channel x time matrix). In order to define PCs that explain a significant proportion of variance in the data, we used non-parametric permutation testing to determine the proportion of variance that can be explained by chance. We randomly shuffled the *F* value time series 1000 times to test the null hypothesis that there is no temporal structure present in the data. Electrodes that exhibited a strong weight (75^th^ percentile) on any of the high variance-explaining PCs as determined by their coefficients were defined as context-encoding. This analytical approach classified electrodes to be context-encoding for 16 patients in PFC (time-locked to HLL; 17 patients showed context-encoding electrodes when time-locked to the behavioral response) and for 11 patients in motor cortex.

#### HFA peak analyses

HFA peak amplitude and timing were estimated on a trial-by-trial basis and used as a proxy of strength and timing of the neural responses, respectively. We included both trials in which participants released the button within the lower and upper limit (correct trials) and trials in which they released the button only after the upper limit (incorrect trials). Amplitude and latencies below the 2.5^th^ or above the 97.5^th^ percentile per channel were considered as outliers and removed from further statistical analysis.

#### HFA single trial regression to behavior

Peak amplitude and latency were computed as described above (see “HFA peak analyses”) and regressed against behavior (RT) via linear regression. We quantified the neuro-behavioral relationship using both full (peak amplitude + latency ~ behavior) and partial linear models (peak amplitude/latency ~ behavior).

#### Estimation of ramping dynamics

To estimate ramping dynamics on a trial-by-trial basis, we quantified the slope of single trial HFA traces using robust linear regression. The slope was estimated from trial start to the HLL.

#### Time-frequency decomposition

We decomposed the raw data into the time-frequency domain using the multitaper method based on discrete prolate spheroidal Slepian sequences in 33 logarithmically spaced bins between 0.5 and 128 Hz. Temporal and spectral smoothing was adjusted to approximately match a 200ms time window and ¼ octave frequency smoothing. To avoid edge artifacts and allow for resolving low frequency activity, decomposition was performed from ±2 sec. surrounding the HLL. As for the HFA analysis (see “HFA extraction”), we normalized the time-frequency data per frequency bin using a bootstrapped baseline distribution (from -0.4 to -0.1s relative to cue onset). Power values were z-transformed according to the means and standard deviations of the bootstrapped distribution. This procedure accounts for the 1/f power drop-off as a function of frequency and minimizes any bias due to baseline differences.

#### Spectral slope estimation

Spectral estimates were obtained by means of a fast Fourier transform (FFT) for linearly spaced frequencies between 1 and 45 Hz after applying a Hanning window and zero padding the data to obtain a fine-grained frequency resolution of 0.25 Hz to improve subsequent background activity estimation. In order to get an estimate of the aperiodic background activity of the power spectrum, we utilized irregular-resampling auto-spectral analysis (IRASA)^32^. IRASA takes advantage of the fact that resampling the original time series by a non-integer resampling factor will leave the 1/f background activity unchanged while systematically shifting the peak frequency at the scale of resampling. Thereby, IRASA disentangles the spectrum into oscillatory (periodic) and 1/f (aperiodic) components. We used the original resampling parameters 1.1 to 1.9 in steps of 0.05^32^ that have also been used in a variety of previous studies^31,54,58^. In a next step, we quantified the spectral slope by means of applying a linear fit to the aperiodic power spectrum in log-log space between 30 to 45 Hz as suggested previously^29^.

#### Time-resolved sample entropy

Sample entropy reflects an information-theoretic measure and captures the complexity of natural time series data^59^. Sample entropy is defined as the negative natural logarithm of the conditional probability that two sequences similar for *m* data points will still match when another data sample (*m* + 1) is added to the sequence:

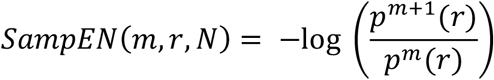

where *m* defines the sequence length, *r* the similarity criterion and defines the tolerance with which two points are considered similar, and *N* the length of the time series to be considered for analysis (*m* = 2 and *r* = 0.2^59,60^). In order to obtain a time series of sample entropy, we estimated sample entropy using a sliding window of 100ms that was shifted in steps of 20ms. Resulting sample entropy time series were smoothed using a 5ms boxcar window to attenuate trial-by-trial variability.

#### HFA peak-triggered average

We conducted a peak-triggered average analysis in order to test 1) whether the HFA is nested into ongoing oscillatory activity, and 2) whether the strength of oscillatory activity is context-dependent. This approach is conceptually similar to spike-triggered averaging used in single unit electrophysiology^61^. Therefore, we detected peaks in the single-trial HFA traces and re-aligned the raw unfiltered data to the detected peak events (segmented ±0.5s surrounding the peaks). To assess the spectral content of the underlying raw traces, we obtained spectral estimates by means of a FFT for linearly spaced frequencies between 1 and 30 Hz after applying a Hanning window and zero padding the data to obtain a frequency resolution of 0.25 Hz. We used IRASA (same parameters and settings as for “spectral slope estimation”) to discount the aperiodic component. Oscillatory residuals were extracted by subtracting the aperiodic spectral component from the original power spectrum.

#### HFA inter-peak-interval

The speed of the HFA traces was quantified by means of computing the interval between two adjacent peaks. We estimated the inter-peak-interval (IPI) on single trials and transformed the distance into frequencies (sampling frequency divided by the time interval between two adjacent peaks). The instantaneous frequency of the HFA amplitude modulation was inferred by the mean of the distribution.

#### Connectivity estimates

We calculated phase-based connectivity metrics between PFC and motor cortex electrodes to infer inter-areal interactions. We first established the presence of undirected phase-based connectivity between PFC and motor cortex by means of the imaginary phase-locking value (iPLV). The iPLV was computed for center frequencies between 3 to 32 Hz (± center frequency/4), logarithmically spaced in steps of 2^1/8^ after band-pass filtering and applying the Hilbert transform^62,63^. Only considering the imaginary part of the phase-locking value removes zero-phase lag contributions^64^. The iPLV was computed as:

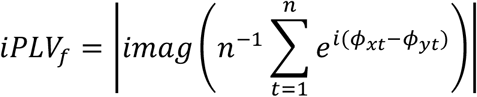

where *n* is the number of time points and *ϕ* reflects the phase angles from electrode *x* and *y* at time *t* and frequency *f*. We first identified the electrode in motor cortex that explained most behavioral variance using linear regression (regressor = HFA timing; response variable = RT). This substantially reduced the degrees of freedom in terms of prefrontal-motor electrode combinations. We then quantified the iPLV between all PFC electrodes and the motor cortex electrode explaining most of the behavioral variance. We have chosen to use behavioral variance explaining electrodes in motor cortex, and not in PFC as motor cortex reflects the final cortical output station to direct behavior^65^. To normalize undirected connectivity, we obtained a permutation distribution by randomly shuffling trial vectors and re-computing the iPLV for every random partition. We further randomly resampled the permutation values 1000 times to approximate a normal distribution. The resulting mean and standard deviations of the bootstrapped permutation distribution were then used to z-normalize the iPLVs. Having established the presence of inter-areal connectivity, we used the phase-slope-index (PSI)^35^ to infer directional connectivity between PFC and motor cortex. We focused our PSI analysis on the low-frequency range given that we observed true oscillatory activity within the low-frequency theta band (**Fig. 4c**). We employed an individualized measure of the PSI using participant-specific peak iPLV frequencies between 2-13 Hz (computed separately per prefrontal-motor electrode pair and using the grand average across all trials) in order to maximize sensitivity and prevent spurious inference on directional prefrontal-motor connectivity^55^. Channel-pairs without a distinct iPLV peak between 2-13 Hz were discarded from the analysis. We computed the PSI between prefrontal-motor electrode pairs on segmented data (zero-padded by 2s on every side) using the corresponding peak iPLV frequency (±3 Hz frequency boundary; linearly spaced). PSI values were z-normalized by means of a permutation distribution that was created by randomly shuffling the frequencies in one vector and recomputing the PSI (1000 iterations)^55^. Note that we used both context-encoding and non-encoding electrodes for undirected and directed connectivity estimates to sample the entire network population.

### Population dynamics

#### Multidimensional distance

The activation state of the full neural population at time *t* can be represented as a point in a *n*-dimensional coordinate system where *n* reflects the number of electrodes (state space). The neural dynamics between the activation state at time *t* and time *t+t*_*n*_ can then be represented as a trajectory through this *n*-dimensional state space^10,16,66^. We quantified the population dynamics by means of the HFA as a proxy for local population activity^17-19^. To investigate whether neural trajectories in the state space are context-dependent, we computed the Euclidean distance between pairwise neural trajectories (e.g. 0% and 75% likelihood of stopping) and then summed the pairwise distances. We used a sliding window of 50ms that was shifted by 20ms in time to obtain a time series of multidimensional distances (**Fig. 5 a/b**). We smoothed the time series using a 25ms boxcar window to attenuate trial-by-trial variability.

#### Euclidean state transitions

We also quantified transitions within neural trajectories separately per context condition (**Fig. 5 a/b**). Thus, we computed the Euclidean distance on single trial trajectories between two adjacent 50ms time windows that were overlapping for 20ms.

#### Dimensionality reduction (PCA)

We used principal component analysis (PCA) to identify linearly uncorrelated population activity patterns and construct a low-dimensional manifold that is embedded in the neural state space spanned by the recorded depth electrodes. We performed PCA on a two-dimensional data matrix (channel x time, trial). The resulting matrix (component x time, trial) was then reshaped into a three-dimensional matrix (trial x component x time) which allowed us to perform single trial analysis in PC space.

#### Identification of coding dimensions

While the top PCs reflect a set of orthogonal dimensions that are optimized to capture maximum variance, they might not always reflect the computationally-relevant subspaces. We used linear discriminant analysis (LDA) to identify the dimensions that carry maximal information about neural dynamics linked to context-integration and movement-execution. We therefore trained two linear classifiers on the PC data. The first classifier was trained to discriminate the type of predictive context, and the second one was trained to discriminate behavioral performance (RT; split into terciles; referred to as action). This procedure allowed us to dissociate neural dynamics linked to the integration of predictive context from subsequent dynamics linked to action. We split the data into training and testing sets using tenfold cross-validation. Because results obtained from cross-validation are stochastic by nature (due to the random assignment of trials into folds), we repeated the analysis five times and then averaged across the repetitions. We applied the LDAs to all PCs in order to identify the dimension that carries most information about our latent variable of interest (note that we only considered PCs that cumulatively explained 99% of the variance and discarded the remaining PCs from the decoding analysis). Decoding traces were then smoothed via application of a 25ms boxcar window. We applied a threshold at chance level to the resulting decoding time series (~33% for both context and action) and set values below chance level to zero. Next, we identified clusters in the decoding time series (adjacent non-zero values) and summed the classification accuracies within each cluster. We defined the PC with the maximum decoding accuracy (largest cluster) as the dimension coding for the latent variable (context or action; referred to as action or context coding dimension). We further created a permutation distribution of classification accuracies by randomly shuffling the trial labels and re-computing the largest cluster from the resulting decoding time series 50 times. We then contrasted the classification accuracies (cluster values) of the identified coding dimension with the generated permutation distribution. We only considered the coding dimension to be valid if the true cluster exceeded the 95^th^ percentile of the permutation distribution. Importantly, we further constrained the dimensions coding for context and action to be orthogonal (distinct PCs). This, however, was empirically the case without adding constraints in 11/14 participants in PFC (*P* = 0.057; two-tailed Binomial test) and 10/10 participants in motor cortex (*P* = 0.002).

#### Determination of oscillatory components in PC space

We obtained spectral estimates for all PCs using IRASA (see “Spectral slope estimation”) for linearly spaced frequencies between 2 and 13 Hz. We then identified the PC with the strongest power in this frequency range.

#### PC-based functional connectivity

We determined the functional connectivity between PCs using power correlations. We computed the correlation coefficient between PC single-trials in PFC and motor cortex. To compare the power correlation across conditions, we normalized the correlation coefficients based on a permutation distribution. We generated the permutation distribution by a random block swapping procedure. This procedure was repeated 1000 times on a trial-by-trial basis to obtain a permutation distribution. Correlation coefficients were then z-transformed using the mean and standard deviation of the permutation distribution.

### Statistical analysis

#### Analysis of variance (ANOVA)

Data were aggregated into ROIs (averaged across electrodes) for statistical testing. We performed a one-way repeated-measures ANOVA using predictive context as a within-subject factor to analyze behavior (**Fig. 1c**), HFA peak latency/amplitude (**Fig. 2a/b**), HFA ramping activity (**Fig. 3a/b**), aperiodic slope (**Fig. 3e/g**), inter-peak interval (**Fig. 4f**) and phase-slope index (**Fig. 4h**). Since not every participant was implanted with electrodes in both PFC and motor cortex, we computed the ANOVA separately for both cortices to estimate the main effect of context onto our latent variable. Significant ANOVA effects were followed by post-hoc testing (two-tailed and corrected for multiple comparisons using the Benjamini-Hochberg procedure^67^). We computed the interaction effect between context x region of interest (PFC, motor cortex) using only a subset of participants that were implanted with electrodes in both regions (N = 11). We considered participant data where z-scores exceeded 3^rd^ standard deviation as outliers.

#### Linear mixed effect models

We confirmed the ANOVA results using linear mixed effect models. Participants were treated as random effects while context and ROI were treated as fixed effects in our model. This approach has been used in previous studies involving human intracranial EEG recordings^55,68^. Model testing was obtained by likelihood ratio tests to compare the models with and without an interaction term (context x ROI). Linear mixed effect models largely confirmed the ANOVA results and are reported in **Supplementary Table 2**.

#### Non-parametric cluster-based-permutation analysis

We used non-parametric cluster-based permutation testing^69^ (as implemented in Fieldtrip^50^) to analyze data in the time (**Fig. 1d**; **2a/b**; **3f/h**; **5a-d, g/h**), frequency (**Fig. 4c/g**) or time-frequency (**Fig. 3c/d**) domain (Monte Carlo method; 10000 iterations; maxsum criterion; two-tailed). Clusters were formed by thresholding a dependent t-test at a critical alpha of 0.05. We generated a permutation distribution by randomly shuffling trial labels and recomputing the cluster statistic. The p-value was then obtained by contrasting the true cluster statistic against the permutation distribution. Clusters were considered to be significant at *P* < 0.05. We also computed interaction effects (context x ROI) using cluster-based permutation testing. We therefore contrasted the difference between two context conditions (75% and 0% likelihood of stop) obtained per ROI using dependent t-tests (only performed on a subset of participants that were implanted with electrodes in both regions). Clusters were considered significant at *P* < 0.05. Note that the cluster-level test statistic reported throughout the text refers to the sum of the F- or t-values in the cluster.

#### Bootstrapping

To control for trial differences across conditions, we used a bootstrap procedure. We randomly resampled as many trials from the two context conditions (0% and 25% likelihood) as there were trials in the 75% condition. This procedure was repeated 500 times, if not stated otherwise. The bootstrapped mean was then considered the final value for the conditions with a higher-trial count^70^.

## Acknowledgements

This work was funded by the Baden Wuerttemberg Foundation (Postdoc Fellowship; RFH), German Research Foundation, Emmy Noether Program (DFG HE8329/2-1; RFH), Hertie Foundation, Network for Excellence in Clinical Neuroscience (RFH), the Research Council of Norway (grant number 240389; AKS, TE, PGL), the Research Council of Norway (Centre of Excellence scheme, grant number 262762; RITMO, RITPART International Partnerships for RITMO Centre of Excellence, grant number 274996; AKS, TE, PGL).

## Author contributions

Conceptualization: TE, AKS, RFH

Methodology: JW, RFH

Investigation: AKS, TE, AOB, AL, IF, SK, PL, JI, RTK

Visualization: JW, RFH

Funding acquisition: AKS, TE, RFH

Project administration: TE, AKS, RFH

Supervision: RFH

Writing – original draft: JW, RFH

Writing – review & editing: AKS, TE

## Data availability

Source data is included as Supporting Information. Raw data are available upon reasonable request from Anne-Kristin Solbakk or Tor Endestad.

## Code availability

Freely available software and algorithms used for analysis are listed where applicable. All code will be made publicly available upon publication on GitHub.

## Additional information

**Supplementary Figure 1.**
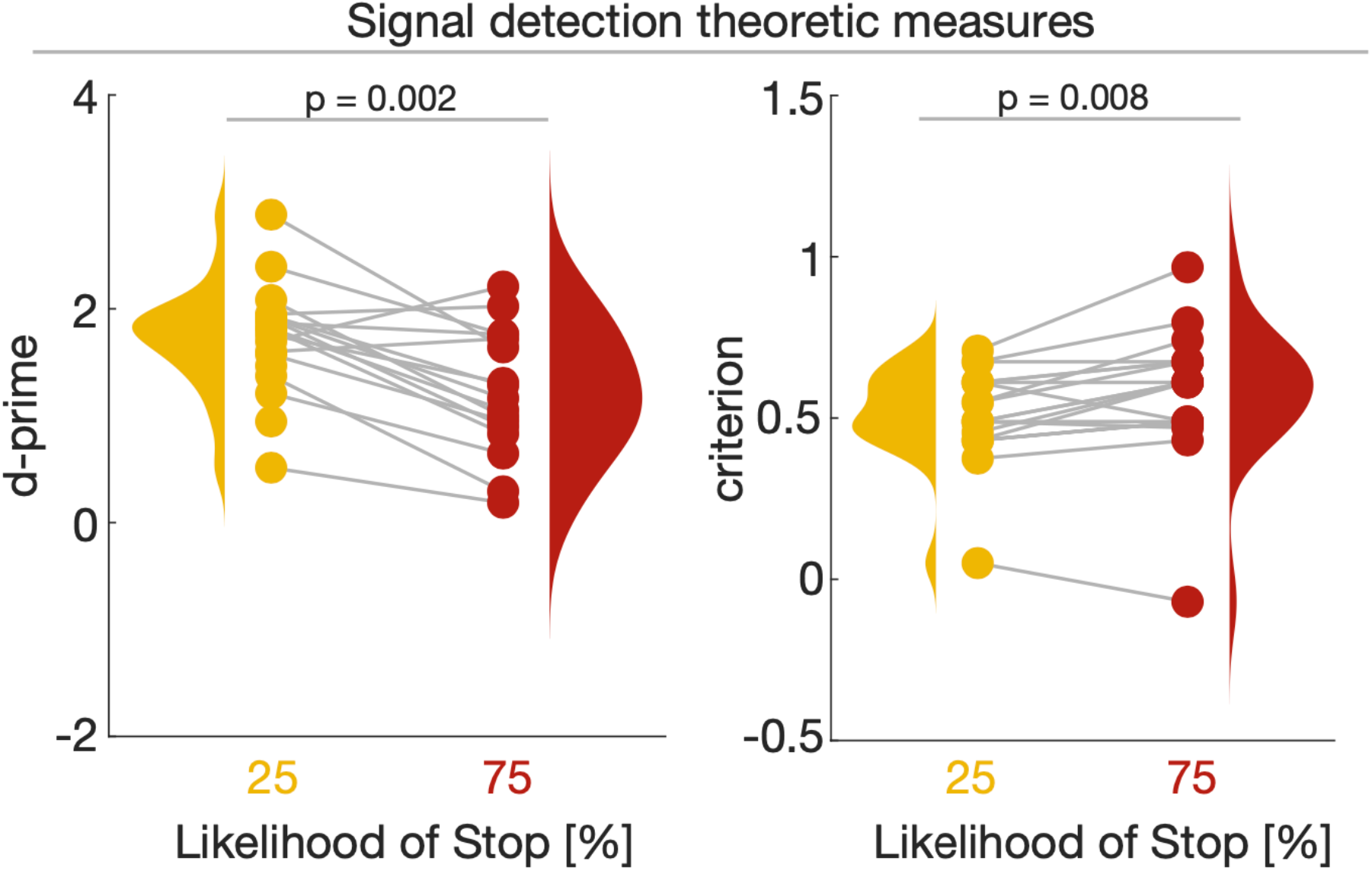
Behavioral uncertainty shifts participant’s sensitivity, but not criterion. Predictive context significantly shifted participant’s sensitivity (d-prime; left panel) and criterion (right panel) indicating a lower discriminability between trial types and an increased tendency to withhold the response with increasing uncertainty.

**Supplementary Figure 2.**
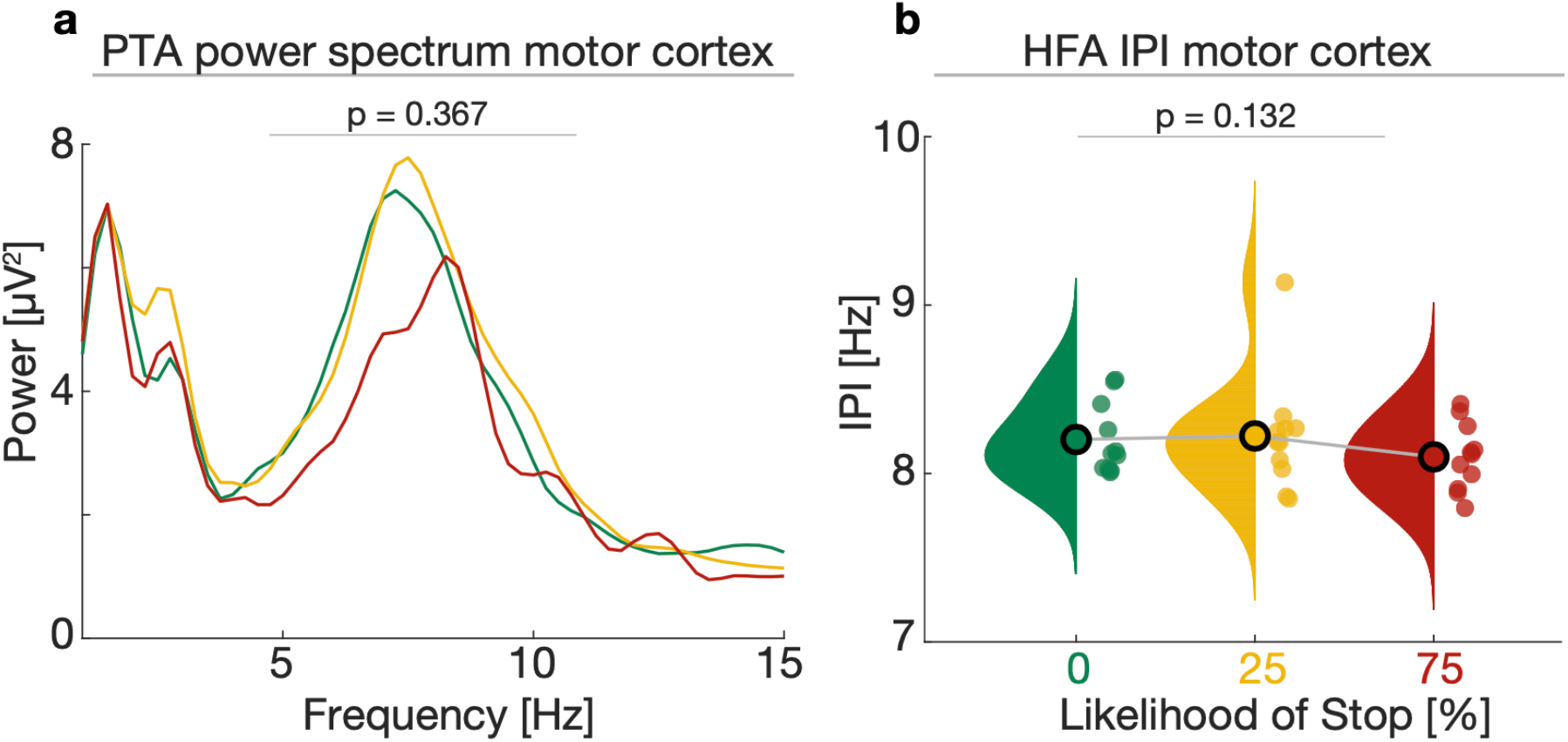
Theta oscillations in motor cortex are not modulated by predictive context. **a**, Grand average 1/f-corrected power spectrum computed on the peak-triggered-average (PTA) time-series using IRASA. The HFA was strongly nested in theta oscillations across all three predictive context conditions as can be seen by the pronounced theta peaks in the power spectrum. **b**, The HFA inter-peak-interval in motor cortex was not significantly modulated by predictive context.

**Supplementary Table 1.** Results for partial linear regression between HFA peak amplitude or peak timing and behavioral response time.

| *PFC* | *df* | *F* | *P* | *R*<sup>2</sup> |
| --- | --- | --- | --- | --- |
| RT ~ Peak Timing | 1,2903 | 61.74 | 5.47 x 10<sup>-15</sup> | 0.021 |
| RT ~ Peak Amplitude | 1,2903 | 30.64 | 3.38 x 10<sup>-8</sup> | 0.01 |

| *Motor Cortex* | *df* | *F* | *P* | *R*<sup>2</sup> |
| --- | --- | --- | --- | --- |
| RT ~ Peak Timing | 1,2017 | 69.51 | 1.38 x 10<sup>-16</sup> | 0.033 |
| RT ~ Peak Amplitude | 1,2017 | 6.88 | 0.008 | 0.0029 |

**Supplementary Table 2.** Linear mixed-effect models supporting the results obtained from the ANOVAs.

| *Modelled* | *Parameter* | *Beta* | *T-statistic* | *df* | *p-value* | *Lower-95 CI* | *Upper-95 CI* | *Random effects (SD)* | *Model comparison (Likelihood ratio / p-value)* |
| --- | --- | --- | --- | --- | --- | --- | --- | --- | --- |
| *HFA peak amplitude (simple model)* | Intercept | 7.416 | 17.921 | 75 | 0 | 6.592 | 8.240 | 1.29 | 0.87 / 0.34<br>(favoring simple model) |
|  | ROI | 0.179 | 1.469 | 75 | 0.146 | -0.064 | 0.423 |  |  |
|  | Prediction | 0.012 | 3.522 | 75 | **< 0.001** | 0.005 | 0.019 |  |  |
| *HFA peak timing (simple model)* | Intercept | -0.103 | -9.014 | 77 | < 0.001 | -0.126 | -0.080 | 0.017 | 0.05 / 0.81<br>(favoring simple model) |
|  | ROI | 0.015 | 3.179 | 77 | **0.002** | 0.005 | 0.024 |  |  |
|  | Prediction | 0.0003 | 2.478 | 77 | **0.015** | < 0.0001 | 0.0006 |  |  |
| *HFA ramping dynamics (simple model)* | Intercept | 0.3 | 0.635 | 77 | 0.526 | -0.639 | 1.240 | 0.934 | 0.37 / 0.54<br>(favoring simple model) |
|  | ROI | 0.029 | 0.158 | 77 | 0.874 | -0.338 | 0.397 |  |  |
|  | Prediction | 0.012 | 2.149 | 77 | **0.034** | 0.0008 | 0.023 |  |  |
| *Aperiodic slope (simple model)* | Intercept | -2.886 | -28.27 | 78 | 0 | -3.089 | -2.683 | 0.271 | 0.38 / 0.53<br>(favoring simple model) |
|  | ROI | 0.111 | 3.232 | 78 | **0.001** | 0.042 | 0.18 |  |  |
|  | Prediction | 0.001 | 0.984 | 78 | 0.328 | -0.001 | 0.003 |  |  |
| *Inter-peak-interval (simple model)* | Intercept | 8.232 | 162.88 | 77 | 0 | 8.131 | 8.332 | 0 | 0.54 / 0.46<br>(favoring simple model) |
|  | ROI | -0.003 | -0.159 | 77 | 0.873 | -0.047 | 0.04 |  |  |
|  | Prediction | -0.001 | -2.561 | 77 | **0.012** | -0.003 | -0.0004 |  |  |
| *PSI* | Intercept | 0.098 | 0.719 | 40 | 0.47 | -0.177 | -0.374 | 0.29 | no other model used |
|  | Prediction | 0.005 | 2.221 | 40 | **0.032** | 0.0004 | 0.009 |  |  |

